# Microsatellites used in forensics are located in regions unusually rich in trait-associated variants

**DOI:** 10.1101/2023.03.07.531629

**Authors:** Vivian Link, Yuómi Jhony A. Zavaleta, Rochelle-Jan Reyes, Linda Ding, Judy Wang, Rori V. Rohlfs, Michael D. Edge

## Abstract

The 20 short tandem repeat (STR) markers of the combined DNA index system (CODIS) are the basis of the vast majority of forensic genetics in the United States. One argument for permissive rules about the collection of CODIS genotypes is that the CODIS markers are thought to contain information relevant to identification only (such as a human fingerprint would), with little information about ancestry or traits. However, in the past 20 years, a quickly growing field has identified hundreds of thousands of genotype-trait associations. Here we conduct a survey of the landscape of such associations surrounding the CODIS loci as compared with non-CODIS STRs. We find that the regions around the CODIS markers are enriched for both known pathogenic variants (>90th percentile) and for SNPs identified as trait-associated in genome-wide association studies (GWAS) (≥95th percentile in 10kb and 100kb flanking regions), compared with other random sets of autosomal tetranucleotide-repeat STRs. Although it is not obvious how much phenotypic information CODIS would need to convey to strain the “DNA fingerprint” analogy, the CODIS markers, considered as a set, are in regions unusually dense with variants with known phenotypic associations.

## Introduction

DNA evidence has played a crucial role in forensic investigations for over three decades (Butler, 2015; Jobling & Gill, 2004; Kayser & de Knijff, 2011; Roewer, 2013). Beginning in the mid-1980s (Gill et al., 1985), forensic practitioners realized that even small numbers of genetic markers—provided that they are sufficiently heterozygous—can provide a nearly unique identifier that rules out the vast majority of people as the source of an unidentified sample. Many governments worldwide began to collect genotypes from highly variable short tandem repeat (STR, also called microsatellite) markers for the purpose of assisting forensic investigations. STR alleles differ from each other by virtue of containing different numbers of repeats of a short (generally 1-6 base pairs) motif sequence (Gymrek, 2017). (STRs of the same length may also differ in their underlying sequence (Gettings et al., 2015), but distinct length classes are the basis for most forensic work.) Because many alleles are possible at an STR locus and STR mutation rates are high, STRs tend to be highly heterozygous (Willems et al., 2014). As a result, small sets of STRs—relatively easily genotyped using technology available in the 1990s—can provide enough information to identify a person from a high-quality single-source DNA sample. Small sets of STRs remain the standard for forensic practice in most countries.

In the United States (US), the Combined DNA Index System (CODIS) markers are the workhorse loci used in forensics. CODIS includes a set of 20 STR markers, 13 of which were established as the original set in the 1990s, and 7 of which were added in 2017 (Hares, 2015). Of the 20 CODIS STRs, 19 are tetranucleotide STRs (i.e. STRs with four-base-pair motifs), and one (D22S1045) is a trinucleotide STR. The X-linked Amelogenin locus is also recorded and may be searched under more restricted circumstances. As of November 2022, CODIS genotypes from 21,791,620 people were accessible to law enforcement via the National DNA Index System (NDIS), and CODIS genotypes had been used as evidence in 622,955 investigations (FBI, 2022).

The broad collection, storage, and use of CODIS genotypes is premised in part on the idea that collection of one’s CODIS genotypes entails only a minimal privacy incursion. When the CODIS markers were expanded from 13 to 20 markers, an explicit goal was to avoid including markers that would allow prediction of disease (Hares, 2012, 2015). The metaphor of a “DNA fingerprint,” sometimes used to describe a person’s CODIS genotypes, conveys this impression, and it has been invoked in legal decisions concerning the CODIS markers, for example the case of *Maryland v. King*, which permitted the collection of CODIS genotypes from arrestees (*Maryland v. King*, 2013).

One piece of evidence that has been marshaled in defense of the claimed phenotypic irrelevance of the CODIS loci is that the CODIS markers themselves have not been associated with known traits. For example, ten years ago, Katsanis & Wagner (2013) scoured the literature and found no record of direct associations between the CODIS markers and any known phenotypes. However, they did note that several of the CODIS markers are intragenic in genes with known phenotypic associations. It is perhaps unreasonable to expect much direct evidence of CODIS-trait associations given that STR markers are seldom tested for association with phenotype directly (but see Wyner et al., 2020). However, our knowledge of phenotypic associations has grown tremendously in the decade since Katsanis & Wagner’s study, prompting a re-examination of their question, in line with calls for systematic reviews of trait information contained in CODIS loci (Kaye, 2014).

Here, we carry out a similar exercise to Katsanis & Wagner, searching widely used genomic databases to characterize the genomic neighborhoods of the CODIS markers. In addition to providing an update to Katsanis & Wagner’s work, we extend it in four main ways. First, we examine the hundreds of thousands of known genotype-phenotype associations identified by genome-wide association study (GWAS) (Buniello et al., 2019; Visscher et al., 2017), particularly those loci near the CODIS markers. Second, we automate most of our procedures, facilitating replication of our work. Third, whereas Katsanis & Wagner considered only very short genomic regions around the CODIS markers (1 kilobase), we consider larger regions as well (10kb and 100kb). Though SNP-STR linkage disequilibrium (LD) tends to be smaller than SNP-SNP LD, SNP-STR LD nonetheless extends over these larger regions (Payseur et al., 2008; Willems et al., 2014), making them relevant for investigation. Finally, Katsanis & Wagner considered only the 13 original CODIS markers and 11 markers suggested for inclusion, seven of which were added in 2017. Here, we consider STR markers across the genome, aggregating data (available as supplementary material) from approximately 1.6 million STRs. We focus our comparisons on 224,092 autosomal tetranucleotide-repeat STRs, as 19 of the 20 CODIS STRs have tetranucleotide repeat motifs.

## Methods

### Data

In January 2023, we downloaded the locations of ~1.6 million STR regions from the hipSTR reference (Willems et al., 2017; http://webstr.ucsd.edu/downloads, direct link https://github.com/HipSTR-Tool/HipSTR-references/blob/master/human/hg19.hipstr_reference.bed.gz). We also downloaded a set of genome-wide annotations from the UCSC Genome Browser (Lee et al., 2020) using the DataIntegrator tool. In particular, we downloaded coding gene locations (Genes and Gene Predictions > NCBI Refseq > RefSeq All and Genes and Gene Predictions > NCBI Refseq > RefSeq Select) from RefSeq (O’Leary et al., 2016), SNP allele frequencies from HapMap (Gibbs et al., 2003) CEU (Variation > HapMap SNPs… > HapMap SNPs CEU), common SNP locations from dbSNP 153 (Sherry et al., 2001) (Variation > dbSNP Archive - dbSNP 153… > Variants), locations of phenotypically relevant variants (Phenotype and Literature > ClinVar Variants… > ClinVar SNVs) from ClinVar (Landrum et al., 2016), trait-associated SNPs discovered in GWAS (Phenotype and Literature > GWAS Catalog) from the GWAS catalog (MacArthur et al., 2017), and the locations of DNase I hypersensitivity clusters (Regulation > ENCODE Regulation - DNase Clusters V3) from ENCODE (Abascal et al., 2020).

All genomic locations were expressed in hg19 / GRCh37 coordinates.

### Data processing

We sought to describe the genomic neighborhoods of all 1.6 million STR regions identified in the hipSTR reference in terms of their density of key annotated features–-in particular, of coding genes, common SNPs, trait-associated variants, and DNase I hypersensitivity sites. Before doing so, we preprocessed the feature data from UCSC in various ways.

For coding gene locations, we used the RefSeq Select set, which contains one entry per curated coding gene (21,432 genes). We also located the transcription start site (TSS) of each gene as either the start or end coordinate of transcription, depending on whether the gene was annotated on the + (TSS = start) or - (TSS = end) strand. To identify SNPs common in people of European ancestries, heavily represented in GWAS (Martin et al., 2019; Popejoy & Fullerton, 2016), we filtered to SNPs with minor allele frequency 1% or larger in the HapMap CEU data, reducing the number of variants from 4,029,798 to 2,705,918. We limited ClinVar variants to those classified as “Pathogenic,” reducing from 1,491,509 variants to 113,412. For DNase I hypersensitivity sites, we limited to sites with the highest signal level (score 1000/1000), reducing the number of sites from 1,949,038 to 160,870.

For the GWAS catalog, we preprocessed in two distinct ways. The GWAS catalog contains one row per unique combination of SNP locus (rsid), study (PubMed ID), and trait, for a total of 392,271 entries. To obtain information about the number of SNPs identified as trait-associated in any GWAS, we first filtered the GWAS catalog to contain only one row per SNP locus, reducing to 183,014 rows. Thus, for counts of numbers of GWAS hits, each SNP rsid counts only once, regardless of how many studies identified it, and regardless of how many traits it was associated with. Next, we sought to identify traits with nearby GWAS associations for each STR. The trait identifiers in the GWAS catalog are not standardized, and many similar traits receive distinct names (for example “HDL cholesterol” and “HDL cholesterol levels” or “Mean corpuscular hemoglobin” and “Mean corpuscular hemoglobin concentration”). To reduce this redundancy and focus on commonly studied traits when counting the number of distinct traits near each STR, we limited to traits with associations reported in at least three distinct studies with the exact same trait name. This reduced the number of traits from 10,399 to 493.

For all features and all STRs, we recorded the distance of the nearest feature to the STR midpoint, and the number of features within 1kb, 10kb, and 100kb of the STR midpoint. For coding gene locations, we kept track of distance to the nearest gene (defined as the distance to the start or end of transcription, whichever is shorter, or 0 if the STR is intragenic) and the nearest TSS separately. For the GWAS catalog, we kept track of the number of GWAS hits within each distance window as well as the number of distinct associated traits (where again, distinctness merely means a non-identical character string). Because of the large size of the dbSNP common variants catalog, we recorded these locations only for the 20 CODIS markers. Additionally, for the CODIS only, we recorded the names of the traits reported as associated in ClinVar and the GWAS catalog, as well as the names of nearby protein-coding genes.

The data processing and analysis scripts, written in R (v. 4.1.2, R Core Team, 2021) and using the data.table package (Dowle et al., 2019), are available at https://github.com/edgepopgen/CODIS_proximity. The output files recording the features proximal to each STR are available in supplementary files.

## Results

### Genetic neighborhoods of the CODIS markers

Table 1 shows the positions of the CODIS markers, the distance to the nearest gene, the names of genes within 100 kilobases (kb) of each marker, and the number of HapMap SNPs at minor allele frequency >1% in the CEU subset of the 1000 Genomes project within 10kb. Half of the 20 CODIS markers are intragenic, as noted previously (Katsanis & Wagner, 2013). Of the remaining 10 markers, 5 have protein-coding genes within 100kb. The CODIS marker with by far the greatest distance to the nearest protein-coding gene in RefSeq Select is D13S317, which is approximately 1.7 megabases (Mb) from the nearest gene. All CODIS markers are within 10kb of several SNPs common in people of European ancestries.

**Table 1.** Locations of the CODIS markers

| marker | chr | Start position<br>(approximate<br>MB, hg19) | Distance to<br>nearest<br>protein-coding gene<br>(0 =<br>intragenic) | Protein-coding<br>genes w/in 100kb, in<br>proximity order | Common SNPs in<br>Hapmap CEU w/in<br>10kb |
| --- | --- | --- | --- | --- | --- |
| D1S1656 | 1 | 230.9 | 0 | *CAPN9, AGT,*<br>*C1orf198, COG2* | 58 |
| TPOX | 2 | 1.5 | 0 | *TPO* | 22 |
| D2S441 | 2 | 68.2 | 29,159 | *C1D* | 22 |
| D2S1338 | 2 | 218.9 | 11,910 | *TNS1, RUFY4* | 11 |
| D3S1358 | 3 | 45.6 | 0 | *LARS2, LIMD1* | 7 |
| FGA | 4 | 155.5 | 0 | *FGA, FGB, FGG,*<br>*PLRG1, DCHS2* | 16 |
| D5S818 | 5 | 123.1 | 158,529 |  | 24 |
| CSF1PO | 5 | 149.5 | 0 | *CSF1R, HMGXB3,*<br>*PDGFRB, TIGD6,*<br>*SLC26A2, CDX1* | 36 |
| D7S820 | 7 | 83.8 | 0 | *SEMA3A* | 19 |
| D8S1179 | 8 | 125.9 | 78,404 | *ZNF572* | 19 |
| D10S1248 | 10 | 131.1 | 172,971 |  | 39 |
| TH01 | 11 | 2.2 | 0 | *TH, INS, IGF2,*<br>*ASCL2* | 21 |
| vWA | 12 | 6.1 | 0 | *VWF, ANO2* | 33 |
| D12S391 | 12 | 12.5 | 28,998 | *MANSC1, LRP6,*<br>*BORCS5* | 32 |
| D13S317 | 13 | 82.7 | 1,729,158 |  | 21 |
| D16S539 | 16 | 86.4 | 157,803 |  | 59 |
| D18S51 | 18 | 60.9 | 0 | *BCL2, KDSR* | 25 |
| D19S433 | 19 | 30.4 | 15,972 | *URI1* | 22 |
| D21S11 | 21 | 20.6 | 778,252 |  | 22 |
| D22S1045 | 22 | 37.5 | 0 | *IL2RB, TMPRSS6,*<br>*C1QTNF6, SSTR3,*<br>*KCTD17, RAC2* | 38 |

Table 2 gives information about pathogenic variants identified in ClinVar and GWAS hits within 10kb of each CODIS marker. Six of the ten intragenic CODIS markers are within 10kb of variants identified as pathogenic in ClinVar, ranging from two variants identified for CSF1PO to 25 for TH01. Sixteen of the 20 CODIS markers are within 10kb of at least one SNP identified as a GWAS hit, with TH01 again recording the most trait-associated nearby variants, with 10. TH01 is intragenic to the tyrosine hydroxylase gene *TH*, which plays an important role in synthesizing dopamine from its amino acid precursor, tyrosine (Nagatsu et al., 2019).

**Table 2.** Phenotypic associations within 10kb of the CODIS markers from ClinVar and the GWAS catalog

| marker | ClinVar variants | ClinVar traits | GWA S hits | GWAS commonly studied traits |
| --- | --- | --- | --- | --- |
| D1S1656 | 0 |  | 0 |  |
| TPOX | 12 | Deficiency of iodide peroxidase;<br>Neurodevelopmental disorder | 2 |  |
| D2S441 | 0 |  | 1 |  |
| D2S1338 | 0 |  | 1 | Height |
| D3S1358 | 0 |  | 0 |  |
| FGA | 22 | Hepatocellular carcinoma;<br>Congenital afibrinogenemia;<br>Familial visceral amyloidosis, Ostertag type;<br>Hypofibrinogenemia; Familial hypodysfibrinogenemia;<br>Familial dysfibrinogenemia;<br>Dysfibrinogenemia; Abnormal bleeding | 4 | Fibrinogen; Height; Ischemic stroke; Stroke; Venous thromboembolism |
| D5S818 | 0 |  | 3 | Amyotrophic lateral sclerosis;<br>Total body bone mineral density |
| CSF1PO | 2 | Brain abnormalities, neurodegeneration, and dysosteosclerosis | 7 | Aspartate aminotransferase levels; Monocyte count; Serum total protein level |
| D7S820 | 0 |  | 1 | Obesity-related traits |
| D8S1179 | 0 |  | 3 | Platelet count |
| D10S1248 | 0 |  | 0 |  |
| TH01 | 25 | Permanent neonatal diabetes mellitus; not specified;<br>Autosomal recessive DOPA responsive dystonia; Inborn genetic diseases; Dystonic disorder | 10 | Cystatin C levels; Height; Hematocrit; Hemoglobin; Hemoglobin concentration; Type 1 diabetes; Type 2 diabetes |
| vWA | 17 | von Willebrand disorder; von Willebrand disease type 3; Abnormality of coagulation; von Willebrand disease type 1 | 1 |  |
| D12S391 | 0 |  | 1 |  |
| D13S317 | 0 |  | 2 | Hippocampal volume |
| D16S539 | 0 |  | 6 | Appendicular lean mass; Optic cup area; Response to statin therapy |
| D18S51 | 0 |  | 2 | Heel bone mineral density |
| D19S433 | 0 |  | 1 |  |
| D21S11 | 0 |  | 0 |  |
| D22S1045 | 4 | Ichthyosis; Immunodeficiency 63 with lymphoproliferation and autoimmunity | 4 | Asthma; Eosinophil counts; Rheumatoid arthritis; Tuberculosis |

### Comparisons with other autosomal tetranucleotide-repeat STRs

To place the properties of the CODIS markers in context, we compared them with the other 224,092 autosomal, tetranucleotide-repeat STRs in the hipSTR reference (Willems et al., 2017). (Although one of the CODIS markers, D22S1045, is a trinucleotide-repeat locus, we focused our comparisons on tetranucleotide-repeat loci.) Figure 1 shows the distribution of the CODIS markers (orange) compared with non-CODIS autosomal tetranucleotide STRs (gray) with respect to their proximity to protein-coding genes, ClinVar pathogenic sites, GWAS hits, unique commonly-studied traits associated with nearby GWAS hits, and DNase I hypersensitivity sites. For four of these feature categories, we show the distance to the nearest feature and the count of features within 1kb, 10kb, and 100kb. For commonly studied GWAS traits, we do not show the distance to the nearest feature. The figures suggest that the CODIS STRs are not systematically less informative about traits than non-CODIS STRs in any category, and in fact, the 10kb and 100kb windows surrounding the CODIS markers appear to harbor more trait-associated variants than average, as identified by ClinVar and the GWAS catalog.

**Figure 1.**
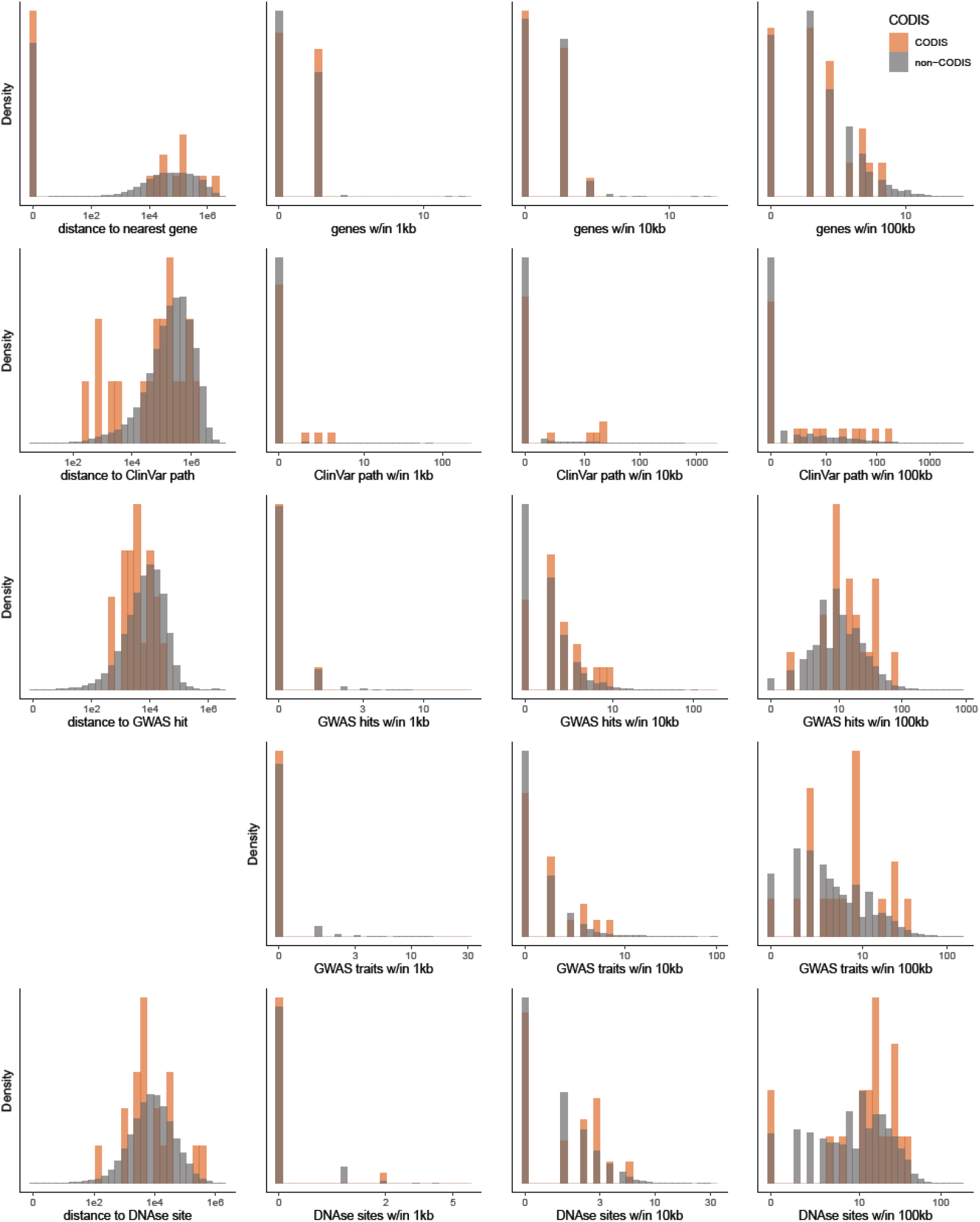
The values of the CODIS loci (orange histogram) compared with non-CODIS autosomal tetranucleotide-repeat STRs (grey) on variables relating to their proximity to phenotype-relevant features. The first column shows distance to the nearest feature, and the second through fourth columns show the number of features within 1kb, 10kb, and 100kb. The rows, in order, show genes included in the RefSeq Select set, variants annotated as pathogenic in ClinVar, SNPs identified as trait-associated in GWAS studies, traits included in at least 3 GWAS studies with associated variants nearby, and DNase I Hypersensitivity sites. The horizontal axes are displayed on a log scale; we added one to all values to avoid taking the logarithm of zero.

Figure 2 shows, for the same features as in Figure 1, the mean of the CODIS markers (dashed orange line) compared with the mean of 10,000 random sets of 20 tetranucleotide markers. The percentiles at which the CODIS average falls on each of these distributions, along with the distributions for TSS and HapMap SNPs common in CEU, are shown in Table 3. Figure 2 and Table 3 confirm the visual impression from Figure 1. The CODIS markers, as a set, are unusually dense with nearby SNPs common in CEU, ClinVar variants marked pathogenic, and GWAS hits. For GWAS hits, the CODIS appear average in their number of hits within 1kb, but above the 90th percentile in the number of hits within 10kb or 100kb. At larger window sizes, the CODIS markers also appear to be in neighborhoods unusually dense in high-scoring DNase I hypersensitivity sites.

**Figure 2.**
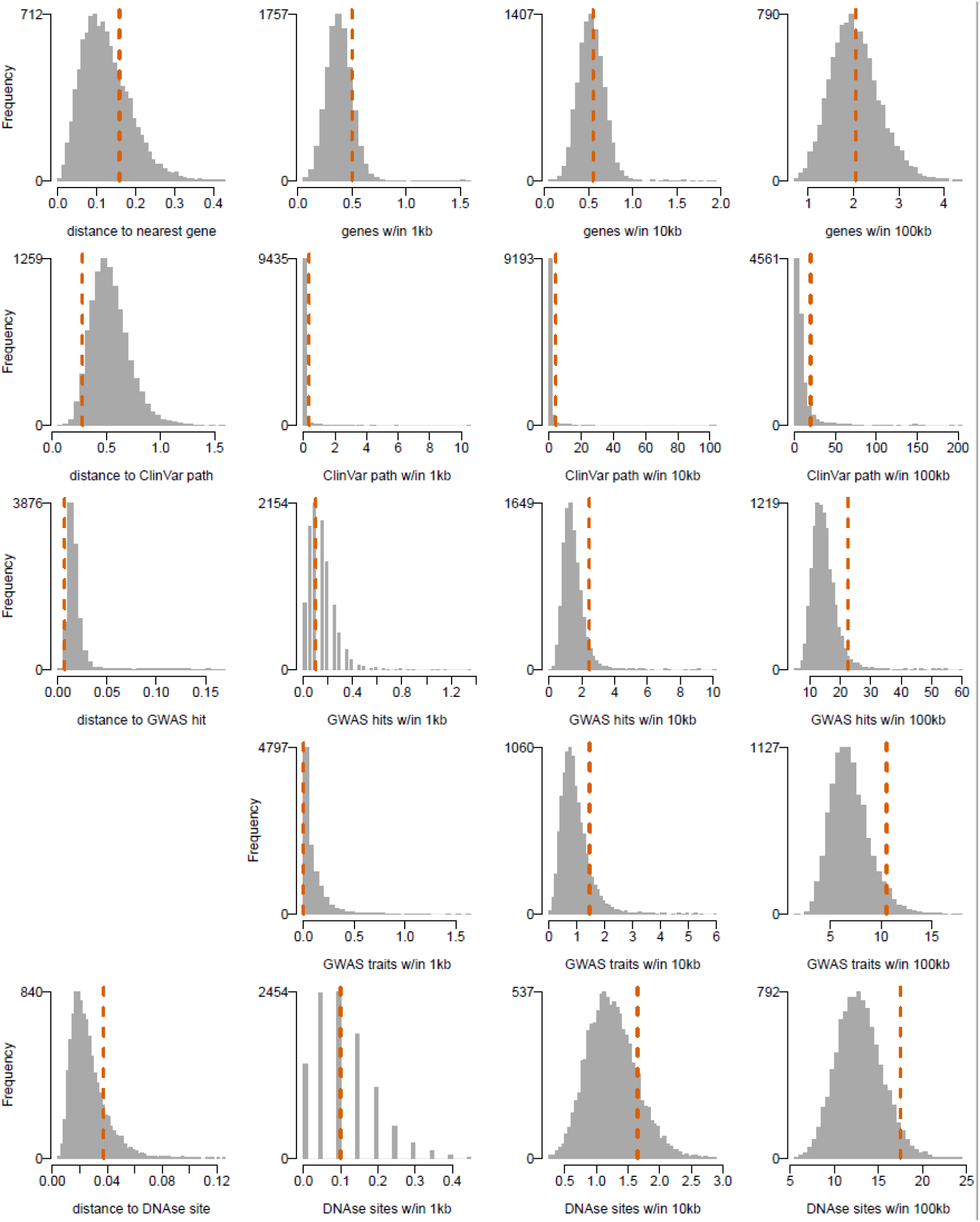
The mean of the 20 CODIS markers (dashed orange line) compared with random sets of 20 non-CODIS autosomal tetranucleotide-repeat loci. The variables shown are the same as in Figure 1.

**Table 3.** Percentiles of the CODIS markers as a set compared with 10,000 random sets of 20 tetranucleotide autosomal STRs

|  | Proximity to nearest* | w/in 1kb | w/in 10kb | w/in 100kb |
| --- | --- | --- | --- | --- |
| RefSeq Select TSS | 50.5 | 96.9 | 77.4 | 67.1 |
| RefSeq Select gene | 26.2 | 86.1 | 57.7 | 54.2 |
| HapMap common SNPs in CEU | 99.9 | 97.2 | 99.7 | 99.0 |
| ClinVar pathogenic variants | 96.1 | 97.0 | 97.4 | 92.2 |
| GWAS hits | 98.9 | 48.6 | 94.7 | 96.7 |
| GWAS well-studied traits | - | 22.7 | 87.7 | 95.6 |
| DNase I Hypersensitivity sites | 16.1 | 62.8 | 85.2 | 96.0 |

\*"Proximity" percentile is 100 minus the "distance" percentile.

Comparing the CODIS markers with sets of random autosomal STRs of irrespective of motif length from one to six (1,527,057 markers in the hipSTR reference) produces results very similar to those obtained for tetranucleotide-repeat STRs (Supplementary Table 1 and Supplementary Figure 1).

We considered whether the unusually high number of GWAS hits and ClinVar pathogenic variants near the CODIS markers might be explained by other features of the CODIS markers. The CODIS markers are 50% intragenic (compared with 39% of non-CODIS tetranucleotide-repeat STRs), and intragenic markers might be expected to be nearer trait-associated variants than intergenic markers. Further, the CODIS markers appear to be in genomic regions with unusually high numbers of SNPs common in people of European ancestry. Since such SNPs are the targets of association in GWAS studies, the high SNP density might explain the high density of GWAS hits.

Table 4 shows Spearman correlations in the non-CODIS autosomal tetranucleotide STRs among intragenic status and the counts of the features in Table 3 (i.e. TSSs, genes, pathogenic variants, GWAS hits and traits, and DNase hypersensitivity sites) within 10kb. (Analogous information for 100kb windows is shown in Supplementary Table 2.) Although intragenic STRs have somewhat more ClinVar pathogenic variants and GWAS hits within 10kb, the correlations between intragenic status and these features are not large (max Spearman rho = 0.22 for ClinVar pathogenic variants). Moreover, comparing the CODIS means to 10,000 random sets of non-CODIS tetranucleotide STRs matched for intragenic frequency (50%) produces a table of percentiles extremely similar to Table 3 (Supplementary Table 3). The correlations between the number of nearby common SNPs and GWAS hits (or ClinVar pathogenic variants) are even smaller than those for intragenic status (Spearman’s rho < 0.1), and in fact, they are mostly negative for counts within 100kb (Supplementary table 1), suggesting that density of nearby SNPs does not explain the unusually high numbers of phenotypic associations near the CODIS markers.

**Table 4.**
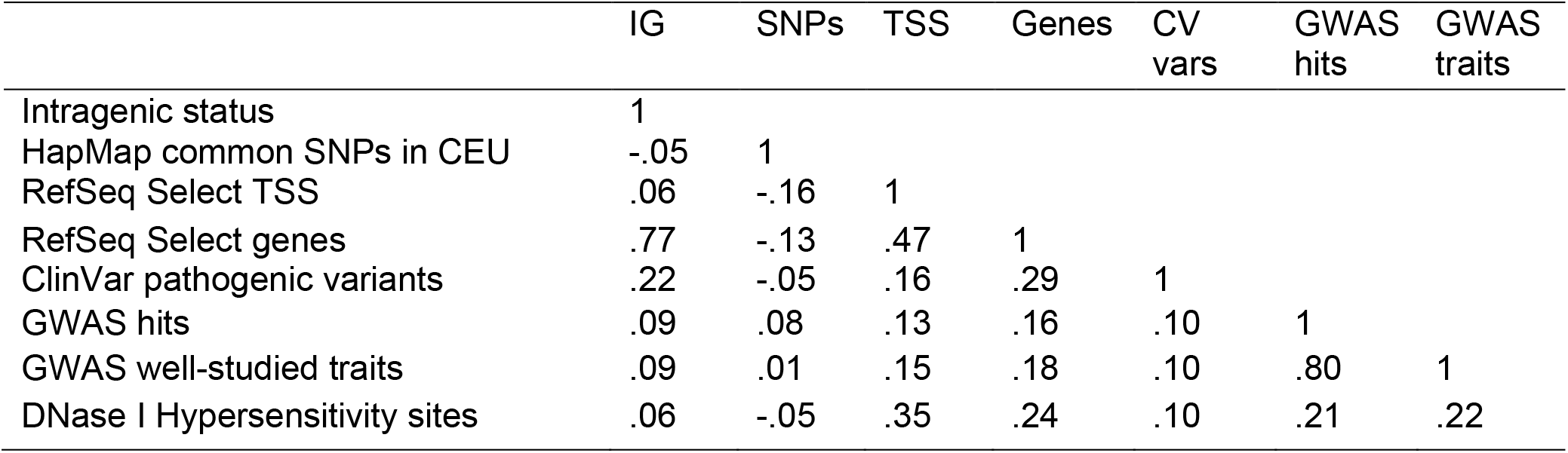
Spearman correlations among key measurements for non-CODIS tetranucleotide STRs (within 10kb)

|  | IG | SNPs | TSS | Genes | CV<br>vars | GWAS<br>hits | GWAS<br>traits |
| --- | --- | --- | --- | --- | --- | --- | --- |
| Intragenic status | 1 |  |  |  |  |  |  |
| HapMap common SNPs in CEU | -.05 | 1 |  |  |  |  |  |
| RefSeq Select TSS | .06 | -.16 | 1 |  |  |  |  |
| RefSeq Select genes | .77 | -.13 | .47 | 1 |  |  |  |
| ClinVar pathogenic variants | .22 | -.05 | .16 | .29 | 1 |  |  |
| GWAS hits | .09 | .08 | .13 | .16 | .10 | 1 |  |
| GWAS well-studied traits | .09 | .01 | .15 | .18 | .10 | .80 | 1 |
| DNase I Hypersensitivity sites | .06 | -.05 | .35 | .24 | .10 | .21 | .22 |

## Discussion

We find that, in comparison with other autosomal tetranucleotide-repeat STRs, the CODIS loci are remarkably rich in nearby variants with known phenotypic associations. The most extreme example is TH01, which has the most known pathogenic variants within 10kb (25) and also the most SNPs within 10kb implicated in GWAS studies (10). Almost 20 years ago, John Butler (2006) wrote that “One core STR locus that has gotten a bad reputation over the years for supposed linkage to genetic diseases is TH01,” going on to note the inconsistent nature of association evidence at the time. Our results are apparently consistent with the reputation TH01 developed among forensic practitioners in the first decade of CODIS’s use. After TH01, the markers with the most known pathogenic variants within 10kb were FGA (22) and vWA (17), and those with the most SNPs identified as trait-associated by GWAS within 10kb were CSF1PO (7) and D16S539 (6).

Although four of these five markers with most evidence of possible trait association (all but D16S539) are intragenic, the unusual proximity of the CODIS to phenotype-associated variants is not explained by the fact that 50% of the CODIS markers are in intragenic regions (compared with 39% of non-CODIS tetranucleotide-repeat STRs). It is also not easily explained by the CODIS markers’ closer proximity to SNPs with minor alleles common in people of European ancestries, since the density of such SNPs is not strongly associated with the presence of either known pathogenic variants or SNPs identified as trait-associated in GWAS.

These results do not constitute direct evidence that the CODIS markers themselves are associated with any phenotypes. However, some degree of correlation (i.e. linkage disequilibrium (LD)) is expected between STRs and SNP markers over these genomic distances (Payseur et al., 2008; Willems et al., 2014). Although the high mutation rates of STRs reduce their LD with surrounding SNPs, genetic drift continually generates LD that is slow to be removed by recombination or nullified by back mutations (Payseur et al., 2008). Direct evidence of whether the CODIS markers (or other STRs) are associated with, or causal for, phenotypes of interest is starting to appear (Gymrek, 2017). We emphasize, however, that from the perspective of phenotype prediction, whether the CODIS markers are causal is not the central concern; any reproducible associations, even if they stem from LD with other causal markers, would still have some predictive utility.

These results add to other lines of evidence suggesting that the CODIS markers are not completely free of phenotypic or other genetic information. For example, the CODIS markers, on closer analysis, turn out to contain substantial ancestry information, despite their low values of F_ST_ (Algee-Hewitt et al., 2016). Further, because the CODIS markers are correlated with—i.e. in LD with—surrounding single nucleotide polymorphism (SNP) markers, it is sometimes possible to identify CODIS and genome-wide SNP genotypes as coming from the same individual, even when the sets of markers in the two datasets are disjoint (Edge et al., 2017; Kim et al., 2018). Most recently, direct examination of the CODIS markers provides suggestive evidence that some of them are associated with gene expression levels in some tissues (Bañuelos et al., 2022).

To be clear, the accuracy of phenotype predictions from the CODIS markers is not expected to be high in absolute terms for most phenotypes. The ability to predict a trait from genotype is limited by the trait’s heritability (Visscher et al., 2008), and for a wide range of complex traits, the best current predictions from genome-wide SNP data are not particularly accurate (Thompson et al., 2022). A small set of STRs will not outperform genome-wide SNPs at phenotype prediction except in rare cases. In general, whether the phenotype predictions developed directly from CODIS represent privacy incursions will depend on at least (a) the standard for how accurate prediction needs to be to be considered a privacy incursion, (b) the number and effect sizes of causal alleles in or near the CODIS markers, and (c) the degree to which a trait is associated with ancestry, which can be noisily reconstructed from CODIS genotypes (Algee-Hewitt et al., 2016). What is clear is that the CODIS markers are not likely to be less informative about phenotypes than other, similar loci. This statement is analogous to the one made by Algee-Hewitt et al. (2016), who found that the CODIS markers are no less informative about ancestry than comparison markers.

It is not clear why the regions around the CODIS markers are unusually dense with phenotypic associations. The GWAS era had not yet begun at the time when the CODIS markers were selected. One possibility is simply bad luck—the original architects of the CODIS system happened to choose sites that would later be identified as near phenotype-associated sites. Another possibility is that there is some other feature or set of features of the CODIS markers that led to both their being considered favorably by the designers of CODIS and that also meant they would be near sites with trait associations, or at least sites that were liable to be discovered as trait-associated. Future work may consider this possibility.

It is not clear why the regions around the CODIS markers are unusually dense with phenotypic associations. The GWAS era had not yet begun at the time when the CODIS markers were selected. One possibility is simply bad luck—the original architects of the CODIS system happened to choose sites that would later be identified as near phenotype-associated sites. Another possibility is that there is some other feature or set of features of the CODIS markers that led to their being considered favorably by the designers of CODIS and that also meant they would be near sites with trait associations, or at least sites that were liable to be discovered as trait-associated. One clue may be the enrichment of high-signal DNase I hypersensitivity sites near the CODIS markers that we observed. DNase I sites are a hallmark of accessible chromatin, and have been relied upon in searches for regulatory elements, including enhancers and promoters (Chen et al., 2018). Chromatin accessibility may also influence the ease of PCR amplification of STRs. Because ease of genotyping by PCR was a factor in the initial selection of the CODIS markers (Butler, 2006), it is possible that the CODIS markers are more likely to be near regulatory elements. Future work may consider this possibility.

In *Maryland v. King* (2013), Justice Kennedy wrote for the majority that the CODIS loci “come from noncoding parts of the DNA that do not reveal the genetic traits of the arrestee.” This statement was part of the majority’s argument that CODIS genotypes can be thought of as a “DNA fingerprint,” a piece of information useful for identification but not informative about any of a person’s traits or medical information. It followed for the majority that collection and storage of CODIS genotypes, like that of fingerprints, is an appropriate part of a routine pre-trial booking procedure. It is not obvious how much information about other traits the CODIS markers would need to convey in order to invalidate the Court’s premise, nor is it yet clear how much information they actually do convey. At the same time, it appears that any attempt to choose markers for CODIS that convey unusually small amounts of information about phenotypes compared with other STRs does not seem to have been successful.

An acknowledgment that CODIS genotypes may be more revealing than previously assumed may prompt rethinking of the patchwork of highly variable local practices governing CODIS genotype collection, storage, and access (Joh, 2015; Murphy & Tong, 2020; Roth, 2019) and influence considerations regarding universal forensic DNA databases (Miller & Smith, 2022). We advocate, along with Kaye (2014), that biomedical literature continue to be monitored in order to ascertain the phenotypic information accessible to a person with access to CODIS profiles (Bañuelos et al., 2022; Wyner et al., 2020). More generally, we advocate that practices surrounding CODIS profiles should be informed by a framework that considers CODIS genotypes not as isolated pieces of information but as components of a genome connected via linkage disequilibrium produced by recombination, mutation, and our shared evolutionary history (Edge et al., 2017; Kim et al., 2018).

### Limitations of the study

This study is limited by ascertainment biases present in the various databases we considered. To take one example, the GWAS catalog is a function of the actual associations identified in GWAS, which means that associations with widely studied traits, with SNPs included in or well imputed by genotyping arrays commonly used for GWAS, and associations that are more easily detectable in people of European ancestries are more likely to be included. Our data processing procedures, which aimed mainly to arrive at simple summaries of high-confidence features, may also have introduced additional ascertainment biases. Another limitation is that we cannot estimate the actual association between STRs and traits, merely the positions of trait-associated variants nearby.

## Supporting information

Supplement_STRproximityInfo

Supplement_CODISproximityInfo

## Acknowledgments

MDE is funded by NIH grant R35 GM137758. Y.J.A.Z. was supported by the NIH Bridges Fellowship (R25-GM048972) and a Genentech Foundation Fellowship. We thank Andy Clark for suggesting chromatin accessibility as a hypothesis for the co-occurrence of CODIS loci and phenotype-associated variants.

## Author contributions

Conceptualization, MDE and RVR; Methodology, MDE and RVR; Software, MDE, YJAZ, and LD; Data Analysis, MDE, VL, YJAZ, R-JR, and JW; Writing - Original Draft, MDE; Writing - Review and Editing VL, YJAZ, R-JR, JW, LD; Visualization, MDE and RVR; Supervision MDE, RVR, and VL.

## Declaration of interests

The authors declare no competing interests.

---

**Figure S1.**
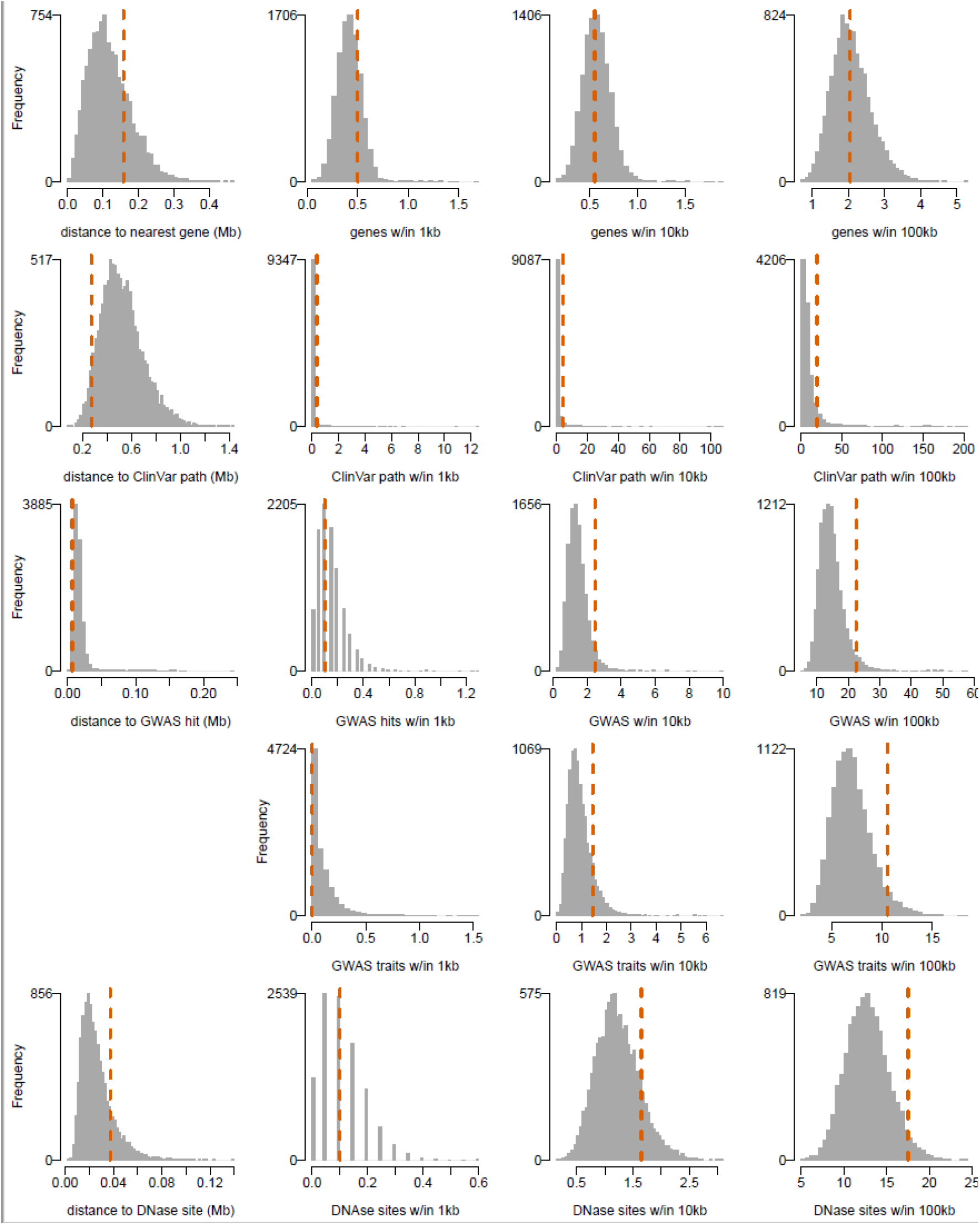
The mean of the 20 CODIS markers (dashed orange line) compared with random sets of 20 non-CODIS autosomal STR loci with repeat lengths from one to six. The variables shown are the same as in Figure 1.

**Supplementary Table 1.**
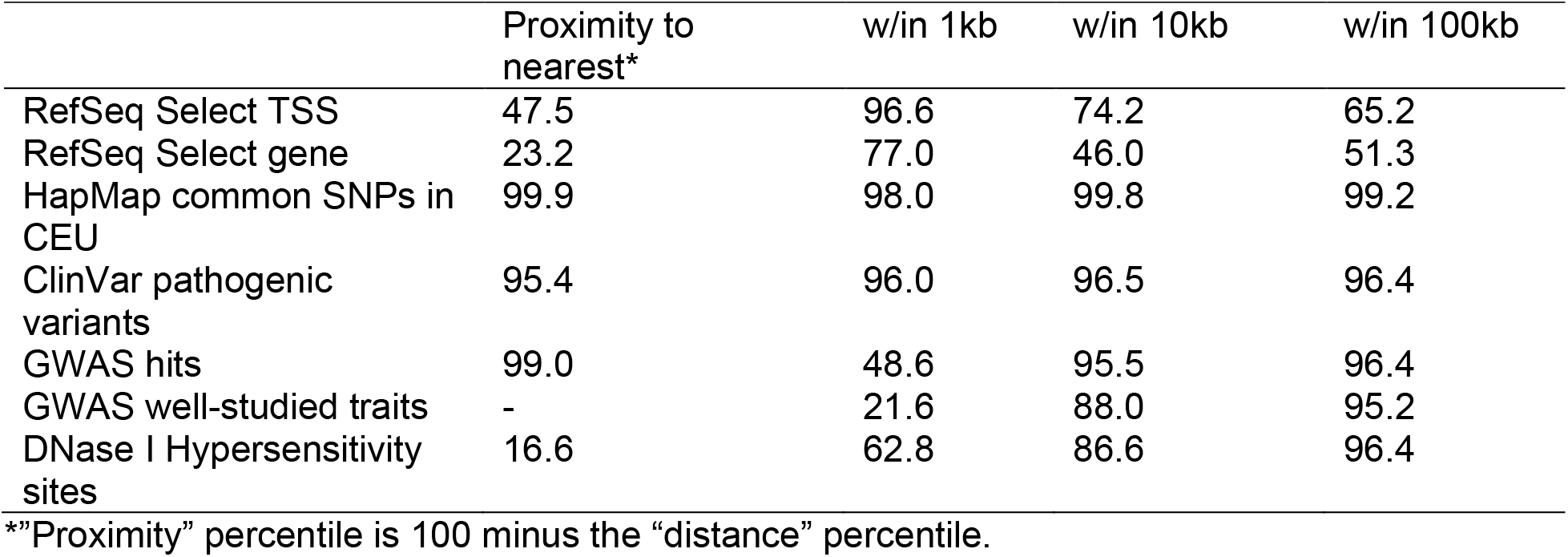
Percentiles of the CODIS markers as a set compared with 10,000 random sets of 20 autosomal STRs with repeat motif lengths ranging from 1-6

|  | Proximity to nearest* | w/in 1kb | w/in 10kb | w/in 100kb |
| --- | --- | --- | --- | --- |
| RefSeq Select TSS | 47.5 | 96.6 | 74.2 | 65.2 |
| RefSeq Select gene | 23.2 | 77.0 | 46.0 | 51.3 |
| HapMap common SNPs in CEU | 99.9 | 98.0 | 99.8 | 99.2 |
| ClinVar pathogenic variants | 95.4 | 96.0 | 96.5 | 96.4 |
| GWAS hits | 99.0 | 48.6 | 95.5 | 96.4 |
| GWAS well-studied traits | - | 21.6 | 88.0 | 95.2 |
| DNase I Hypersensitivity sites | 16.6 | 62.8 | 86.6 | 96.4 |

\*"Proximity" percentile is 100 minus the "distance" percentile.

**Supplementary Table 2.**
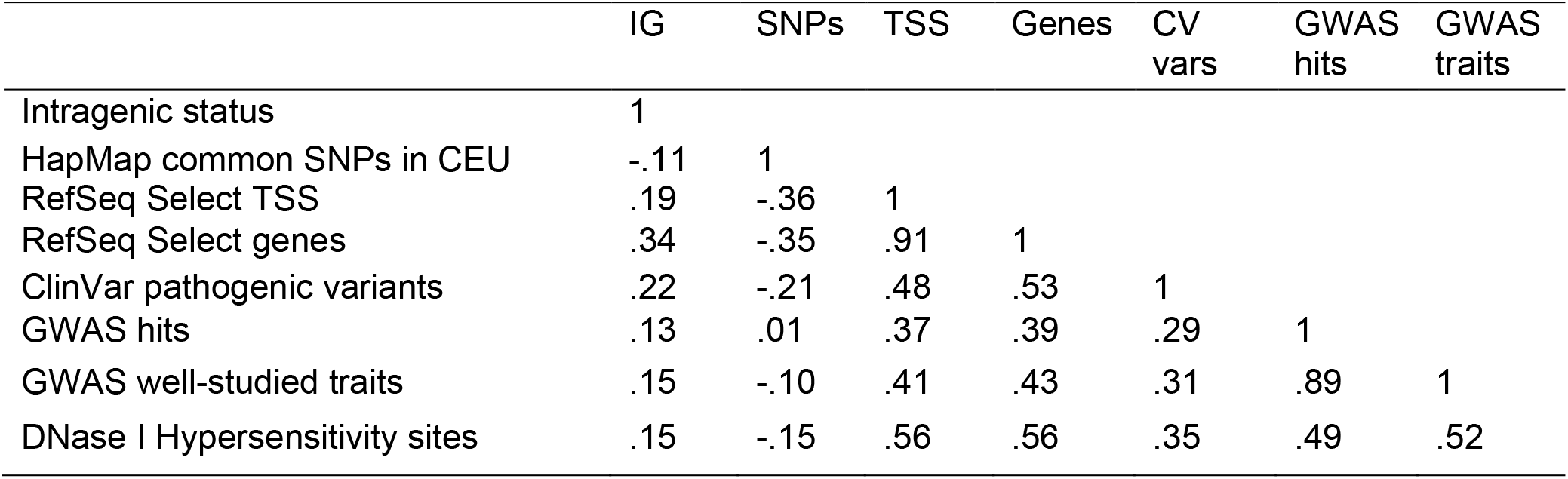
Spearman correlations among key measurements for non-CODIS tetranucleotide STRs (within 100kb)

|  | IG | SNPs | TSS | Genes | CV<br>vars | GWAS<br>hits | GWAS<br>traits |
| --- | --- | --- | --- | --- | --- | --- | --- |
| Intragenic status | 1 |  |  |  |  |  |  |
| HapMap common SNPs in CEU | -.11 | 1 |  |  |  |  |  |
| RefSeq Select TSS | .19 | -.36 | 1 |  |  |  |  |
| RefSeq Select genes | .34 | -.35 | .91 | 1 |  |  |  |
| ClinVar pathogenic variants | .22 | -.21 | .48 | .53 | 1 |  |  |
| GWAS hits | .13 | .01 | .37 | .39 | .29 | 1 |  |
| GWAS well-studied traits | .15 | -.10 | .41 | .43 | .31 | .89 | 1 |
| DNase I Hypersensitivity sites | .15 | -.15 | .56 | .56 | .35 | .49 | .52 |

**Supplementary Table 3.**
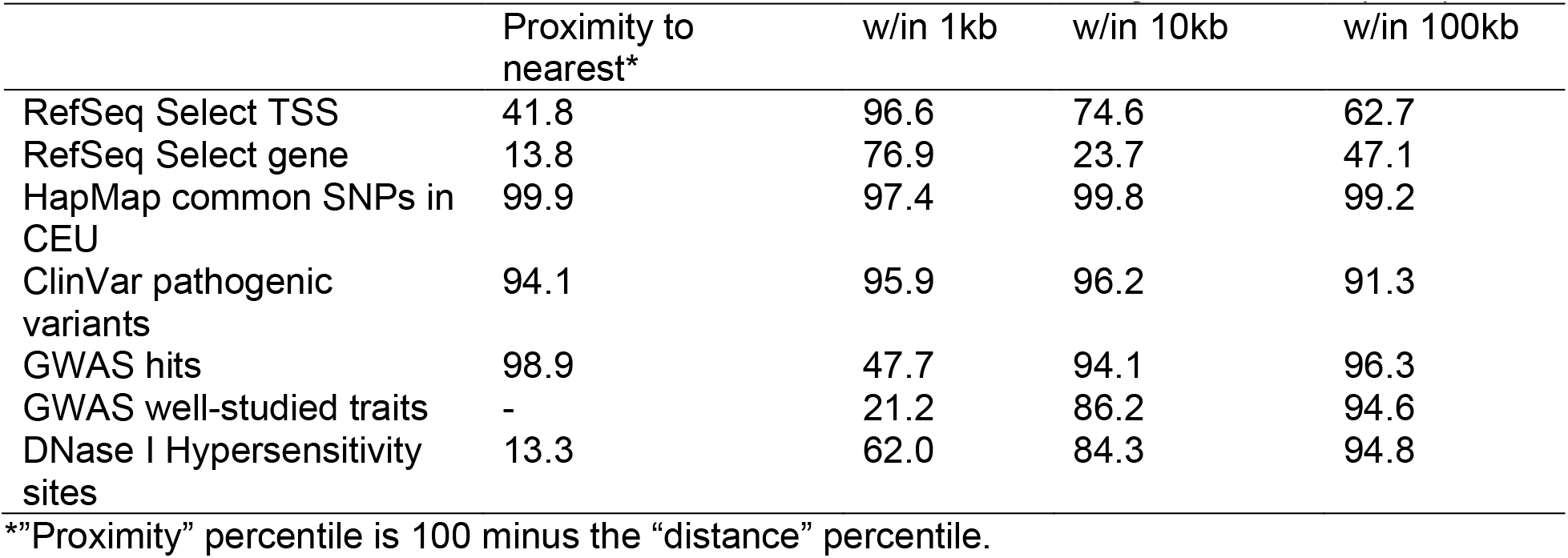
Percentiles of the CODIS markers as a set compared with 10,000 random sets of 20 tetranucleotide autosomal STRs, matched for intragenic fraction (50%)

|  | Proximity to<br>nearest* | w/in 1kb | w/in 10kb | w/in 100kb |
| --- | --- | --- | --- | --- |
| RefSeq Select TSS | 41.8 | 96.6 | 74.6 | 62.7 |
| RefSeq Select gene | 13.8 | 76.9 | 23.7 | 47.1 |
| HapMap common SNPs in<br>CEU | 99.9 | 97.4 | 99.8 | 99.2 |
| ClinVar pathogenic<br>variants | 94.1 | 95.9 | 96.2 | 91.3 |
| GWAS hits | 98.9 | 47.7 | 94.1 | 96.3 |
| GWAS well-studied traits | - | 21.2 | 86.2 | 94.6 |
| DNase I Hypersensitivity<br>sites | 13.3 | 62.0 | 84.3 | 94.8 |

\*"Proximity" percentile is 100 minus the "distance" percentile.

## Notes

https://github.com/edgepopgen/CODIS_proximity

### Competing Interest Statement

The authors have declared no competing interest.

